# An inducible BRCA1 expression system with *in vivo* applicability uncovers activity of the combination of ATR and PARP inhibitors to overcome therapy resistance

**DOI:** 10.1101/2025.08.05.668495

**Authors:** Elsa Irving, Alaide Morcavallo, Jekaterina Vohhodina-Tretjakova, Anna L. Beckett, Michael P. Jacques, Rachel S. Evans, Jennifer I. Moss, Anna D. Staniszewska, Josep V. Forment

## Abstract

Poly(ADP-ribose) polymerase inhibitors (PARPi) have transformed cancer therapy for patients harbouring homologous recombination repair (HRR) deficiencies, notably BRCA1/2 mutations. However, resistance to PARPi remains a clinical challenge, with restoration of BRCA1 function via hypomorphic variants representing an understudied scenario. Here, we engineered a doxycycline-inducible BRCA1 expression system in the BRCA1-mutant, triple-negative breast cancer cell line MDAMB436, permitting controlled analysis of functionally distinct BRCA1 hypomorphs *in vitro and in vivo*. Among multiple BRCA1 variants generated—including RING, coiled-coil, and BRCT domain mutants—only overexpression of the Δexon11 hypomorph robustly conferred resistance to olaparib and carboplatin, with drug sensitivity correlating to Δexon11 expression levels. While Δexon11 BRCA1 mediated HRR restoration, its efficiency was consistently lower than full-length BRCA1, as measured by RAD51 foci formation and interaction with repair partners such as PALB2. *In vivo*, tumours expressing Δexon11 BRCA1 exhibited only partial resistance to olaparib compared to those expressing full-length BRCA1. Importantly, the combination of olaparib and the ATR inhibitor, ceralasertib, overcame Δexon11-mediated resistance, impairing RAD51 foci formation in Δexon11-expressing cells. Our findings identify a dose-dependent, hypomorphic HRR restoration by Δexon11 BRCA1, help explain the variable resistance observed in BRCA1-mutant pre-clinical models expressing this hypomorph and propose ATR inhibition in combination with PARPi as a clinical strategy to counteract therapeutic resistance mediated by Δexon11 BRCA1 hypomorphs.

**Statement of significance:** This work demonstrates that ATR inhibition can overcome PARP inhibitor resistance mediated by BRCA1 Δexon 11 hypomorphs, supporting combination therapy as a promising strategy for PARPi-resistant BRCA1-mutant cancers.

## Introduction

Inhibitors of poly(ADP ribose) polymerases (PARPi) have revolutionized the treatment landscape for cancer patients with tumours harbouring mutations and/or genetic signatures of homologous recombination repair (HRR) defects, including those affecting the *BRCA1* or *BRCA2* tumour suppressor genes. Accordingly, several PARPi are now approved as monotherapies and in combinations for patients with tumours carrying such alterations in different disease settings. However, PARPi resistance arises, with several molecular mechanisms driving resistance having been described in clinical and pre-clinical settings (1).

BRCA1 is a multi-functional protein, with several domains involved in different aspects of DNA repair and cell cycle regulation (**Figure 1A**) (2). The N-terminal RING domain is required for its interaction with BARD1 and is essential for the enzymatic ubiquitin E3 ligase activity of the BRCA1-BARD1 complex (3). The central region of the protein is encoded in a single exon (exon 11), and it is a largely unstructured domain that contains a binding region for the RAD51 recombinase, among other features (4). The coiled-coil domain that sits immediately downstream to the exon 11 sequence is required for the interaction of BRCA1 with the PALB2-BRCA2-RAD51 complex, which is essential for the role of BRCA1 in HRR (5). And the C-terminal BRCT domains of BRCA1 mediate protein-protein interactions with BRIP1, CTIP and the BRCA1-A complex, among others (2). Interestingly, pathogenic mutations in BRCA1 span the entire length of the protein, suggesting that all domains could be important for its function as a tumour suppressor protein (6). However, whether all its functions are required to confer resistance to platinum drugs or PARPi is a matter of debate. As such, expression of hypomorphic versions of the BRCA1 protein has been associated with resistance to platinum drugs and PARPi in pre-clinical models. Hypomorphs lacking part or the entire N-terminal RING domain of BRCA1 have been described *in vitro, in vivo* and in patient-derived xenograft (PDX) models, and have been postulated to be produced by alternative transcriptional START sites downstream of the sequence encoding the RING domain in the BRCA1 transcript (7-10). Similarly, alternative splicing isoforms of BRCA1 have been described that bypass most or the entire exon 11 sequence (BRCA1 Δ11 or Δ11q) (11), which are invariable detected *in vitro* (12) and in PDX models (13) where the original BRCA1 pathogenic mutation lies within the exon 11 of the gene. Hypomorphs lacking one or both BRCT domains of BRCA1 have also been described *in vitro* and in PDX models, and their expression has been linked to chaperone-mediated stabilisation (14) and genomic rearrangements (15), and associated with resistance to treatment (10).

**Figure 1.**
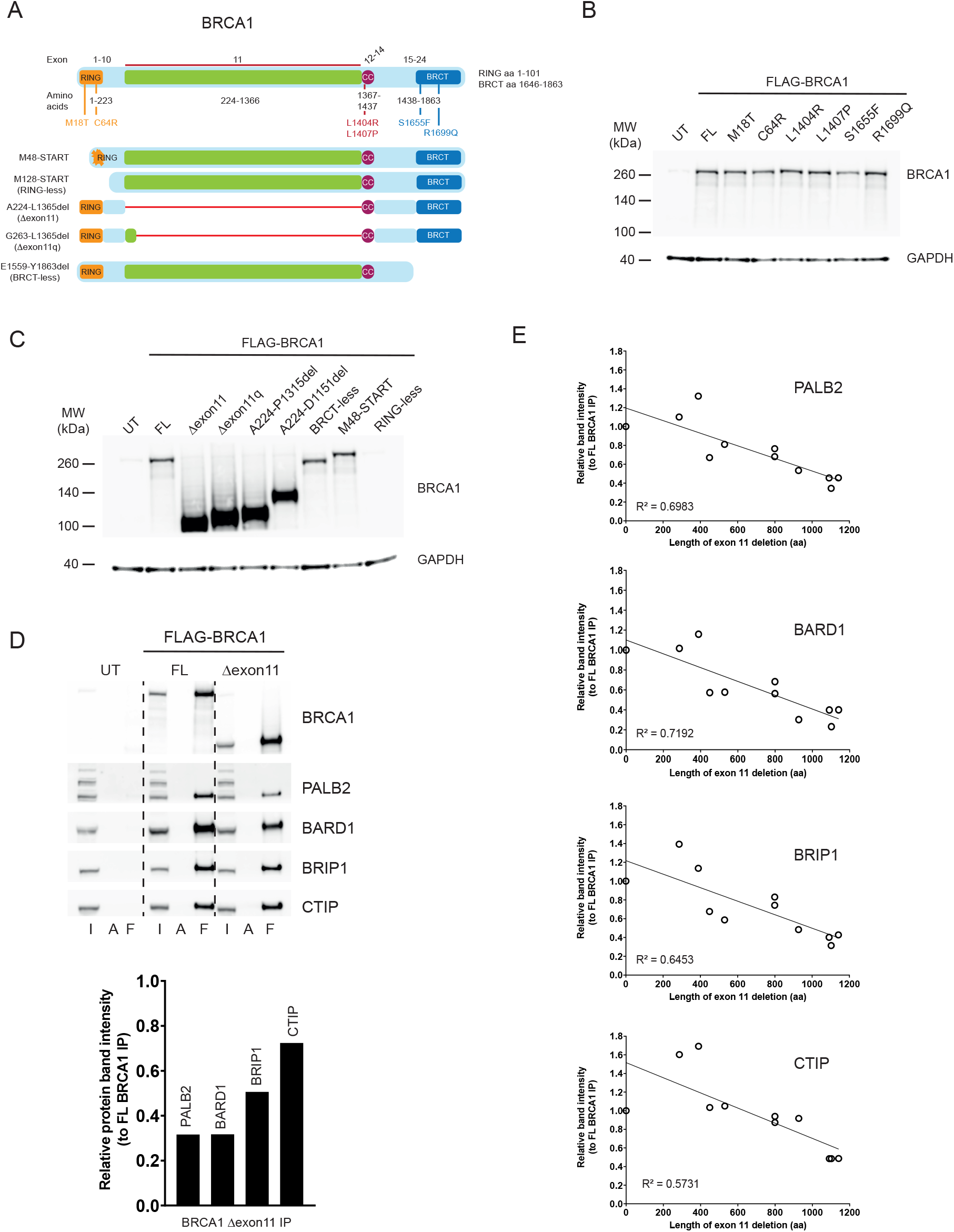
**A**) Schematic of the BRCA1 protein and its functional domains, including the single amino acid changes analysed in this study (top). Deletions representing hypomorphic variants affecting different domains are depicted at the bottom. **B**) Western blot showing expression in HEK293T cells of FLAG-tagged BRCA1 constructs containing the full length (FL) protein or the different amino acid changes assessed in this study. GAPDH was used as loading control. UT: untransfected. **C**) Western blot showing expression in HEK293T cells of FLAG-tagged BRCA1 constructs containing the full length (FL) protein or the different deletion mutants assessed in this study. GAPDH was used as loading control. UT: untransfected. **D**) *Top panel*: co-immunoprecipitation experiments in HEK293T cells with full length (FL) or Δexon11 FLAG-BRCA1. *Bottom panel:* relative pull-down efficiency of BRCA1 interactors with the FLAG-BRCA1 Δexon11 construct, compared to FL protein. I: input; A: pull down with protein A beads; F: pull down with FLAG beads. **E**) Relative pull-down efficiency of BRCA1 interactors with an allelic series of FLAG-BRCA1 deletions inside exon 11, compared to FL protein.

Modelling the effect of BRCA1 hypomorph expression in pre-clinical models has been difficult due to two major problems: the detrimental effect of clinically relevant point mutations on BRCA1 protein levels (5,16) and the lack of *in vivo* inducible expression systems to assess the impact of varying levels of hypomorph expression on therapy responses. To circumvent these issues, we describe here the development of a doxycycline-inducible BRCA1 expression system in the clinically relevant BRCA1 mutant, triple-negative breast cancer cell line MDAMB436. We further demonstrate the utility of this system by describing the requirement of substantial levels of overexpression of the BRCA1 hypomorph lacking the entire exon 11 of the gene (BRCA1 Δexon11) to generate resistance *in vivo* to PARPi, and by identifying the combination of PARPi with inhibitors of the DNA-damage response (DDR) kinase ATR as a way to overcome such resistance.

## Results

### BRCA1 hypomorphic versions display different protein-protein interactions

To better understand the importance of the different BRCA1 functions and domains with respect to response to PARPi, we decided to generate an allelic series capturing point mutations and deletions in different BRCA1 regions (**Fig 1A**). Regarding mutations, we produced M18T and C64R in the RING domain, as well as S1655F and R1699Q in the BRCT domains, as these are well-described germline pathogenic mutations in BRCA1 that have been shown not to result in substantially reduced protein levels when exogenously expressed in human cells (5,16). In addition, we generated two mutations in the coiled-coil domain (L1404R and L1407P) that have been described to disrupt the interaction between BRCA1 and PALB2, which is essential for HRR (5). Regarding deletions, we generated two variants with alternative START codon sites that partially (M48 START) or completely (M128 START, which we name RING-less) remove the N-terminal RING domain of BRCA1, and that have been postulated as alternative translation start points to generate RING-defective BRCA1 hypomorphs (8). We also produced variants lacking the entire exon 11 of the gene (A224-L1365del, which we name Δexon11) or only retaining its first 40 amino acids (G263-L1365del, which we name Δexon11q), as these can be produced through alternative splicing of the *BRCA1* transcript (11). Finally, we also generated a BRCA1 version lacking the C-terminal BRCT domains (E1559-Y1863del, which we name BRCT-less), as similar BRCA1 hypomorphic variants have been linked to PARPi resistance in preclinical models (14,15).

FLAG-tagged constructs with all these variants were produced and expression was tested upon human cell line transfection. All constructs carrying point mutations were expressed at similar levels to the full length (FL) BRCA1 protein (**Fig 1B**). Deletion constructs in the RING (M48 START) and BRCT regions were also expressed at similar levels compared to the FL, while exon 11 deletions resulted in significantly higher protein levels than the FL protein, as previously observed (12,17) (**Fig 1C**). To assess whether point mutations and deletions affected protein-protein interactions as previously described, we immunoprecipitated the different FLAG-tagged BRCA1 constructs and probed for the RING-interacting protein BARD1, the coiled-coil interacting protein PALB2 and the BRCT interacting proteins BRIP1 and CTIP (5). As expected, RING mutants and deletions (M18T, C64R, M48 START, M128 START) all failed to interact with BARD1, the coiled-coil mutants L1404R and L1407P failed to pull down PALB2 and the BRCT mutants and deletion (S1655F, R1699Q, E1559-Y1863del) did not show an interaction with BRIP1 or CTIP (**Supp Fig S1A**). Despite not being able to detect the BRCA1 RING-less mutant by western blotting (due to the antibody epitope being in the RING domain; **Fig 1C**), we did observe it pulling down PALB2, BRIP1 and CTIP, confirming its expression (M128 START pull downs; **Supp Fig S1A**).

We found that, despite their significantly higher expression, the exon 11 deletions did not seem to co-immunoprecipitate more of the BRCA1 interacting partners (**Supp Fig S1A**). To confirm this observation, we adjusted BRCA1 protein concentration levels between FL and Δexon11 (**Supp Fig S1B**) and quantified the amount of BRCA1 interactors co-immunoprecipitated. This highlighted that the BRCA1 Δexon11 protein, despite being able to co-immunoprecipitate all the interactors tested, it did so less efficiently than the FL protein, particularly in the case of PALB2 and BARD1 (**Fig 1D**). To better understand whether there were any specific sequence requirements inside the region encoded by exon 11 to sustain effective protein-protein interactions, we generated an additional allelic series of deletions of different size inside exon 11, including two (R762-D1151del and E427-S713del) reported as secondary reversion mutations detected in patients progressing on PARPi treatment (**Supp Fig S1C;** (18,19)). Importantly, and even though all of them co-immunoprecipitated the different interaction partners tested (**Supp Fig S1D**), there was a clear correlation between the size of the exon 11 deletion and the ability of BRCA1 to perform such interactions (**Fig 1E**).

### Generation of a doxycycline inducible BRCA1 expression system

To better control the effect on the cellular response to PARPi of the different mutations or deletions introduced in BRCA1, we decided to generate a doxycycline-inducible expression system in the triple-negative breast cancer cell line MDAMB436. We chose this cell line because it carries a hemizygous BRCA1 mutation (5396 + 1G>A mutation in the splice donor site of exon 20) (20) that results in strongly reduced levels of BRCA1 protein (12) and exquisite sensitivity to PARPi *in vitro* and *in vivo* (21). Interestingly, examination of the MDAMB436 genetic make-up in the Cancer Cell Line Encyclopaedia (22) highlighted that, in addition to the described *BRCA1* mutation, this cell line also carries a homozygous deletion of the HRR gene RAD51B (**Supp Fig S2A**). As RAD51B loss confers a certain degree of PARPi sensitivity (23), we first produced a MDAMB436 derivative cell line with constitutive expression of *RAD51B* (MDAMB436-B; **Supp Fig S2B**). We then introduced the tetracycline repressor in both MDAMB436 and MDAMB436-B (**Supp Fig S2C**) and named these cell lines MDAMB436-TR and MDAMB436-B-TR, respectively. We transduced them with constructs expressing LacZ (as control) or FL FLAG-tagged BRCA1, and inducibility of the system was confirmed by western blotting (**Supp Fig S2D**).

To understand the importance of RAD51B expression in the MDAMB436 complementation system, we assessed olaparib sensitivity in MDAMB436-TR (lacking RAD51B expression) and MDAMB436-B-TR (expressing RAD51B) cells complemented with FL BRCA1 or control (LacZ) in the presence of doxycycline induction. As shown in **Fig 2A**, cells expressing RAD51B but not BRCA1 (MDAMB436-B-TR + LacZ) were only marginally more resistant to the PARPi, olaparib (half-maximal inhibitory concentration (IC50) of 14.2 nM), than their *RAD51B* null counterpart (MDAMB436-TR + LacZ cells, IC50 = 4.3 nM), as expected given the dominant effect of BRCA1 loss in driving sensitivity to PARPi. Perhaps surprisingly, however, BRCA1 expression in the absence of RAD51B complementation (MDAMB436-TR + BRCA1 cells) only resulted in mild but significantly increased resistance to olaparib (IC50 = 56.2 nM) when compared to their BRCA1 mutant counterpart (MDAMB436-TR + LacZ cells; IC50 = 4.3 nM). Substantial resistance to olaparib was only achieved by expression of both RAD51B and BRCA1 (MDAMB436-B-TR + BRCA1; IC50 = 3899 nM). Consequently, all further experiments were performed in the MDAMB436-B-TR background (hereafter referred to as BTR cells).

**Figure 2.**
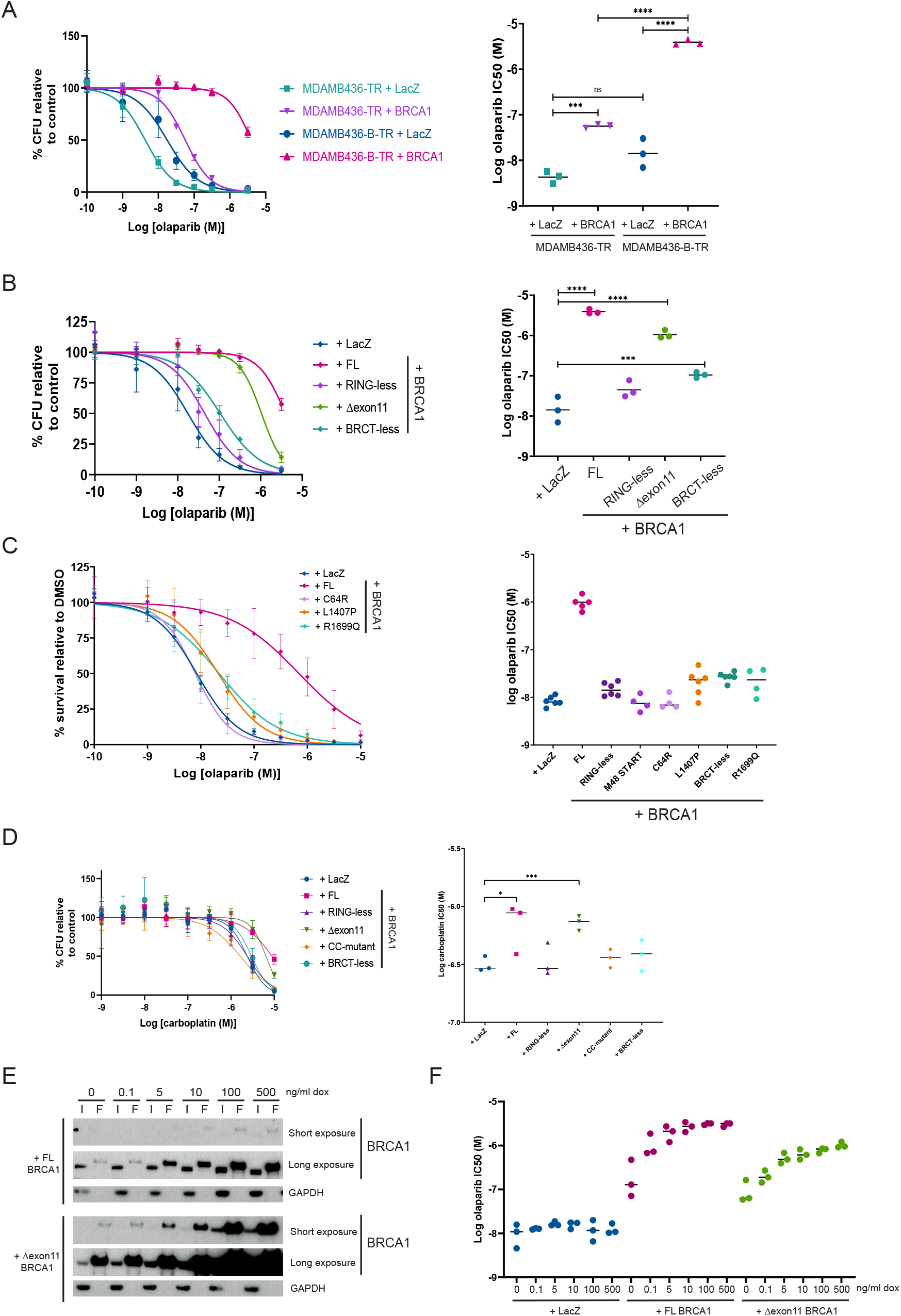
**A**) *Left panel:* dose-response curves of olaparib in colony formation assays in tetracycline-repressor (TR) expressing MDAMB436 cells with or without RAD51B complementation (MDAMB436-B-TR and MDAMB436-TR, respectively) and with (+ BRCA1) or without (+ LacZ) BRCA1 complementation. *Right panel:* Logarithmic half-maximal inhibitory concentration (LogIC50) of olaparib for each cell line. **B**) *Left panel:* dose-response curves of olaparib in colony formation assays in MDAMB436-B-TR cells expressing full-length BRCA1 (+FL), no BRCA1 (+LacZ) or different BRCA1 hypomorphs (+RING-less, +Δexon11, +BRCT-less). *Right panel:* Logarithmic half-maximal inhibitory concentration (LogIC50) of olaparib for each cell line. **C**) *Left panel:* dose-response curves of olaparib in survival assays in MDAMB436-B-TR cells expressing full-length BRCA1 (+FL), no BRCA1 (+LacZ) or different BRCA1 hypomorphs (+C64R, +L1407P, +R1699Q). *Right panel*: Logarithmic half-maximal inhibitory concentration (LogIC50) of olaparib for each cell line. Data for BRCA1 hypomorphs with deletions in the RING (RING-less, M48 START) or BRCT (BRCT-less) domains are included for comparative purposes. **D**) *Left panel:* dose-response curves of carboplatin in survival assays in MDAMB436-B-TR cells expressing full-length BRCA1 (+FL), no BRCA1 (+LacZ) or different BRCA1 hypomorphs (+RING-less, +Δexon11, +BRCT-less, +CC-mutant). *Right panel*: Logarithmic half-maximal inhibitory concentration (LogIC50) of carboplatin for each cell line. **E**) Western blot of immunoprecipitation experiments in MDAMB436-B-TR cells expressing full length (+FL) or Δexon11 BRCA1 and exposed to different doxycycline doses. GAPDH was used as loading control. I = input; F = FLAG immunoprecipitation. **F**) Logarithmic half-maximal inhibitory concentration (LogIC50) of olaparib at the different doses of doxycycline used. All data are from at least 3 biological replicates. Statistical analysis performed using One-Way ANOVA with Holm-Sidak multiple comparisons, ^*^ p<0.05, ^**^ p<0.01, ^***^ p<0.001, ^****^ p<0.0001.

As we wanted to understand how stringent repression of BRCA1 expression was in BTR cells, we tested their sensitivity to olaparib in the presence or absence of doxycycline induction. We observed a mild increase in resistance to olaparib in BTR cells complemented with FL BRCA1 even in the absence of doxycycline treatment (BTR + BRCA1 - dox; IC50 = 109 nM) when compared to non-BRCA1 complemented cells (BTR + LacZ + dox; IC50 = 17 nM) (**Supp Fig S2E**). Inspection of BRCA1 expression by western blotting detected low levels of the FL protein even in the absence of doxycycline induction, potentially explaining the observed increased resistance to olaparib (**Supp Fig S2F**). As expected, doxycycline-induced *BRCA1* expression resulted in significantly increased BRCA1 protein levels (**Supp Fig S2F**) and increased resistance to olaparib (BTR + BRCA1 + dox; IC50 = 3962 nM) (**Supp Fig S2E**).

Taken together, these results show functionality of the doxycycline-inducible BRCA1 complementation system, and the requirement of both RAD51B and BRCA1 complementation in MDAMB436 cells to generate resistance to olaparib to levels comparable to cell lines with no HRR deficiency (23).

### Expression of the BRCA1 Δexon11 hypomorph generates resistance to olaparib and carboplatin in vitro

In addition to the FL BRCA1, BTR cell lines expressing RING mutants (RING-less and C64R), the exon 11 deletion (Δexon11), a coiled-coil mutation (L1407P) or BRCT mutants (BRCT-less and R1699Q) of BRCA1 were also produced (**Supp Fig S2F**). Interestingly, while expression of RING-less (IC50 = 45 nM) or BRCT-less BRCA1 (IC50 = 106 nM) resulted in a non-significant or mild increase in olaparib resistance, respectively, compared to the LacZ control (IC50 = 17 nM), expression of the Δexon11 hypomorph resulted in a substantial increase in olaparib resistance (IC50 = 1047 nM), only surpassed by expression of FL BRCA1 (IC50 = 3962 nM) (**Fig 2B**). Point mutations in the RING (C64R) or BRCT (R1699Q) domains, or a shorter deletion in the RING domain (M48 START), which recapitulate the same protein-binding defects of the more extreme deletions (**Supp Fig S1A**), as well as the coiled-coil mutation L1407P, also resulted in no to mild increase in resistance to olaparib in our system (**Fig 2C**). Similar results were obtained in response to carboplatin treatment, where only expression of FL BRCA1 or the Δexon11 hypomorph generated resistance. Importantly, and in contrast to the olaparib responses, IC50 values for carboplatin between BTR cells complemented with FL BRCA1 or the Δexon11 hypomorph were very similar (**Fig 2D**).

Given that the BRCA1 Δexon11 hypomorph is expressed at significantly higher levels than the FL BRCA1 protein in our system (**Supp Fig S2F**), we performed a doxycycline dose-titration experiment in these cell lines. While expression of both FL and Δexon11 BRCA1 reached saturation conditions at around 100 ng/ml doxycycline (**Fig 2E**), maximal resistance to olaparib was already achieved between 5-10 ng/ml dox in the case of FL BRCA1, and between 100-500 ng/ml dox in the case of the BRCA1 Δexon11 hypomorph (**Fig 2F**). Collectively, these results show that, among all BRCA1 hypomorphs tested, only expression of the Δexon11 protein results in resistance to olaparib and carboplatin to levels associated with restored HRR. Importantly, however, this is only partially achieved at expression levels significantly higher than those of the FL BRCA1 protein, confirming the hypomorphic nature of the Δexon11 deletion.

### Expression of FL BRCA1 or the BRCA1 Δexon11 hypomorph protein generates resistance to olaparib in vivo

We decided to assess whether the increased resistance to olaparib in the BTR complementation system observed *in vitro* could be translated *in vivo*. Interestingly, BTR cells complemented with FL BRCA1 implanted as mouse xenografts developed as tumours at a faster rate than parental, BRCA1 mutant MDAMB436 cells (**Fig 3A**). Analysis of mRNA levels of FLAG-tagged *BRCA1* confirmed its expression in the BTR-derived tumours (**Fig 3B**). As described before (21), tumours generated after implantation of the parental MDAMB436 cell line showed exquisite sensitivity to olaparib in a dose-dependent manner, with the top dose of olaparib used (the clinically relevant 100 mg/kg dose) causing complete tumour regressions (**Fig 3C**). Importantly, xenografts of the BTR cell line complemented with FL BRCA1 displayed strong resistance to olaparib, in agreement with our *in vitro* data, with the 100 mg/kg dose showing a degree of tumour growth inhibition (41% TGI) compared to vehicle control (**Fig 3D**). Also correlating with the *in vitro* data, xenografts of the BTR cell line expressing the BRCA1 Δexon11 hypomorph showed increased resistance to olaparib, but to a lesser extent than their FL BRCA1 complemented counterparts (54% TGI at 100 mg/kg; **Fig 3E**). As expected, expression of FLAG-tagged Δexon11 *BRCA1* transcript was significantly higher than that of the FL transcript (**Fig 3F**).

**Figure 3.**
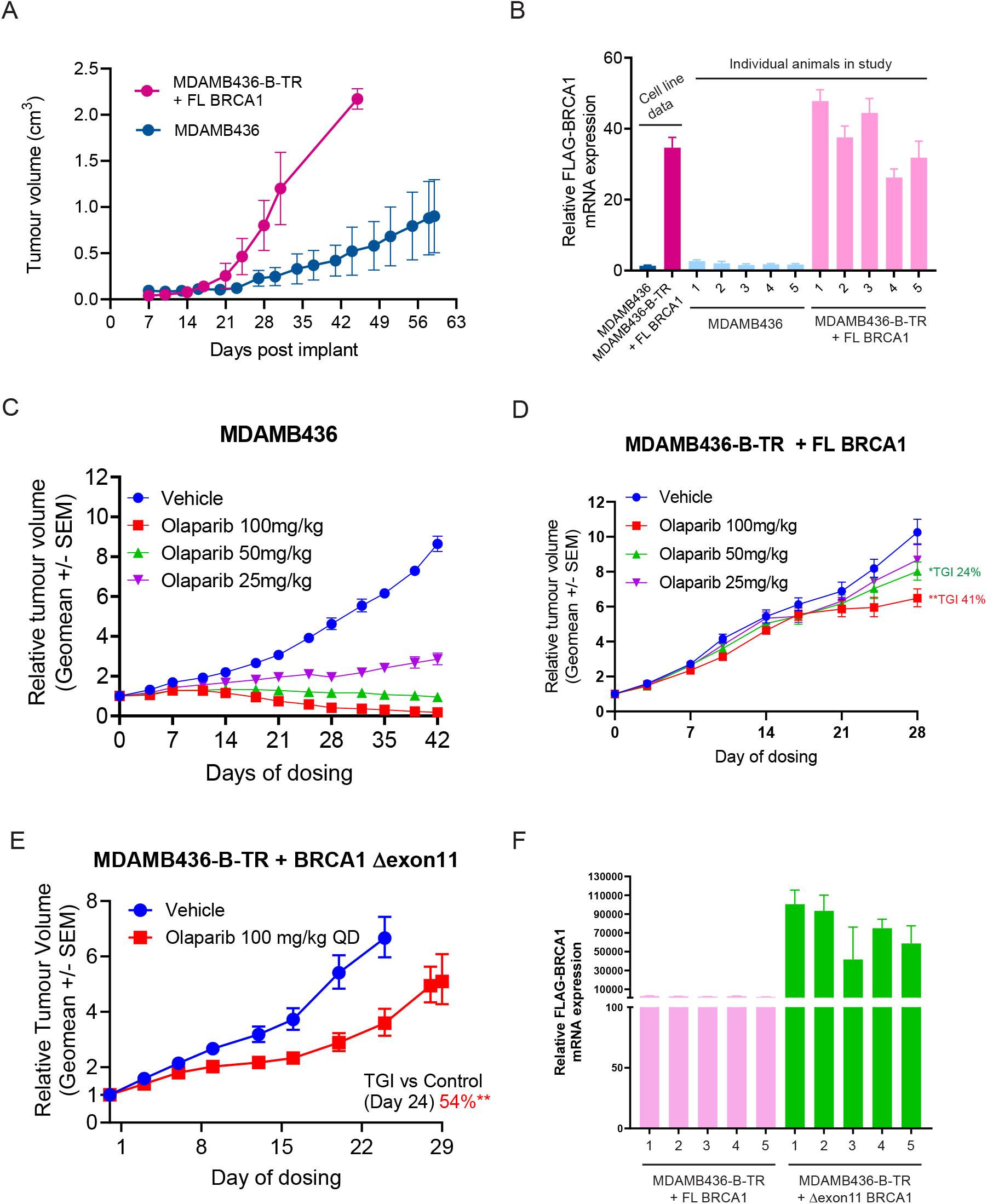
**A**) Tumour volume growth curves of MDAMB436 (BRCA1 mutant) cells or MDAMB436-B-TR cells expressing full length (+FL) BRCA1 in SCID mice. **B**) Relative *FLAG-BRCA1* mRNA expression in tumour samples from individual mice from the experiment in A. Expression is compared to samples taken from *in vitro* culture of the same cell lines and is normalized to that of *GAPDH*. **C**) Dose-response efficacy of olaparib in the MDAMB436 (BRCA1m) xenograft model. **D**) Dose-response efficacy of olaparib in the MDAMB436-B-TR + full length (FL) BRCA1 expressing xenograft model. **E**) Dose-response efficacy of olaparib in the MDAMB436-B-TR + Δexon11 BRCA1 expressing xenograft model. Graphs depict geometrical mean (geomean) of tumour volume ± SEM and percentage TGI (tumour growth inhibition). Statistical significance was evaluated compared to the vehicle group using one-tailed t test (mice n = 8/group). Statistical significance is indicated as follows: ^*^, *P* ≤ 0.05; ^**^, *P* ≤ 0.01; ^***^, *P* ≤ 0.001. F) Relative *FLAG-BRCA1* mRNA expression in tumour samples from individual mice implanted with MDAMB436-BTR cells expressing the full length (FL) or Δexon11 *BRCA1* constructs. Expression is normalized to that of *GAPDH*.

Taken together, these data show that the BTR complementation approach is a useful *in vivo* system to model acquired resistance to olaparib caused by expression of BRCA1 protein variants.

### Combination of olaparib and the ATR inhibitor, ceralasertib, increases efficacy in the BTR BRCA1 Δexon11 model system

Combinations of PARPi with other DDR-targeted agents, as well as with DNA-damaging chemotherapies, are being pursued as a way to overcome PARPi resistance in the clinic (24). Having established an *in vivo* system to understand BRCA1 hypomorph mediated PARPi resistance, we next explored therapeutic approaches to overcome it. With that goal, we assessed combination activity of olaparib with the ATR inhibitor, ceralasertib (25), the WEE1 inhibitor, adavosertib (26), the DNA crosslinking agent, carboplatin and the DNA topoisomerase I inhibitor, SN-38, in BTR cells expressing FL BRCA1 or the BRCA1 Δexon11 hypomorph. As controls, we compared the activity of such combinations in non-complemented BTR cells (LacZ). While combination activity between olaparib and adavosertib, carboplatin or SN-38 was very similar between LacZ, FL or Δexon11 expressing BTR cell lines, as assessed by the highest single agent (HSA) model (27), the ceralasertib combination provided higher score values in BRCA1 Δexon11 expressing cells (**Fig 4A**).

**Figure 4.**
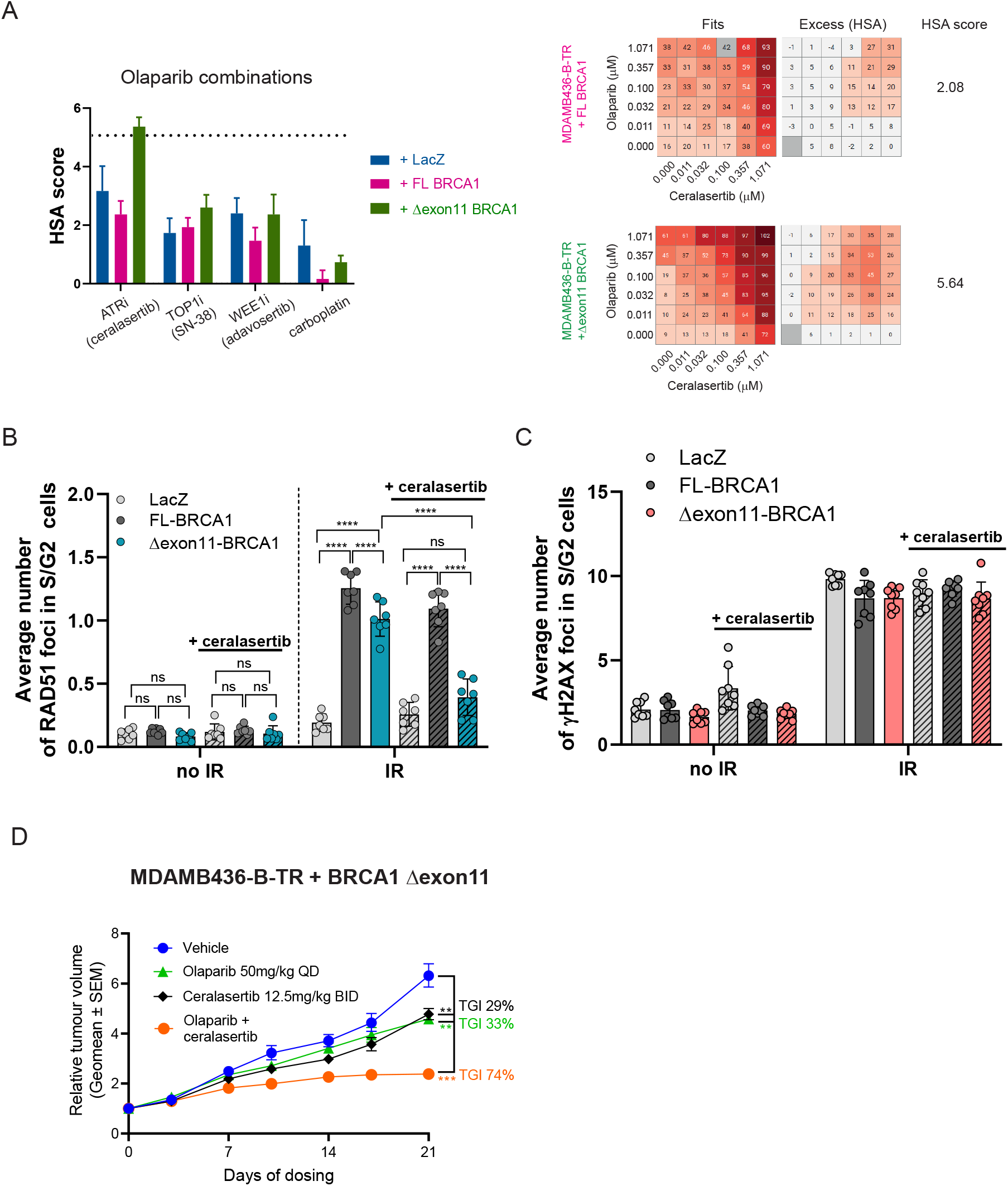
**A**). *Left panel:* combination activity between olaparib and ceralasertib, SN-38, adavosertib or carboplatin in MDAMB436-B-TR cells expressing full length (+FL), Δexon11 or no BRCA1 (+LacZ) protein. Scores above 5 (marked by the dotted line on the graph) are described as synergistic based on the highest single agent (HSA) model. *Right panel*: representative examples of 6×6 combination matrices between olaparib and ceralasertib in MDAMB436-B-TR cells expressing full length (+FL) BRCA1 (top) or Δexon11 BRCA1 (bottom). In the Fits panels, values between 0-100 represent cytostatic effects. The excess (HSA) panels represent the values used to calculate synergy scores. Values coloured in pink/brown represent positive interactions. **B**) Quantification of RAD51 foci formation in MDAMB436-B-TR cells expressing full length (+FL), Δexon11 or no BRCA1 (+LacZ) protein, irradiated (IR) or not with 5 Gy and treated or not with ceralasertib (0.3 µM for 24 hours prior to IR). Statistical analysis was performed using One-Way ANOVA with Sidak multiple comparisons, n=3 biological replicates with 2-3 technical replicates in each experiment; ns – not significant, ^*^ p<0.05, ^**^ p<0.01, ^***^ p<0.001, ^****^ p<0.0001. **C**) Quantification of γH2AX foci formation in MDAMB436-B-TR cells expressing full length (+FL), Δexon11 or no BRCA1 (+LacZ) protein, irradiated (IR) or not with 5 Gy and treated or not with ceralasertib (0.3 µM for 24 hours). Statistical analysis was performed using One-Way ANOVA with Sidak multiple comparisons, n=3 biological replicates with 2-3 technical replicates in each experiment, ns – not significant, ^*^ p<0.05, ^**^ p<0.01, ^***^ p<0.001, ^****^ p<0.0001. **D**) Efficacy of olaparib, ceralasertib or their combination in the MDAMB436-B-TR + Δexon11 BRCA1 expressing xenograft model. Graph depicts geometrical mean (geomean) of tumour volume ± SEM and percentage TGI (tumour growth inhibition). Statistical significance was evaluated compared to the vehicle group using one-tailed t test (mice n = 10/group). Statistical significance is indicated as follows: ^*^, *P* ≤ 0.05; ^**^, *P* ≤ 0.01; ^***^, *P* ≤ 0.001.

Resistance to PARPi in BRCA1 mutant cell lines and tumours caused by acquired expression of BRCA1 hypomorphic or FL proteins is associated with restored HRR, as measured by the ability of cells to form RAD51 foci in response to DNA damage (1,10). We compared the ability of non-complemented (LacZ), FL and Δexon11 BRCA1 BTR cells to form RAD51 foci in response to ionizing radiation (IR) and observed that while LacZ cells failed to show recruitment of RAD51 to DNA damage sites, both FL and Δexon11 BRCA1 BTR cells effectively formed RAD51 foci (**Fig 4B**). Interestingly, Δexon11 BRCA1 BTR cells showed a small but significant defect in RAD51 foci formation when compared to FL BRCA1 complemented cells (**Fig 4B**), indicative of suboptimal HRR proficiency and reflective of the less pronounced resistance of Δexon11 BRCA1 BTR cells to olaparib when compared to FL BRCA1 complemented cells, both *in vitro* and *in vivo* (**Fig 2B and 3D-E**). Strikingly, treatment with the ATRi, ceralasertib, strongly reduced the ability of Δexon11 BRCA1 BTR cells to form RAD51 foci in response to IR, while the effect was much more modest in cells complemented with FL BRCA1 (**Fig 4B**). Reduced levels of RAD51 foci were not due to decreased DNA damage formation, as assessed by comparable levels of the DNA damage induced Ser-139 phosphorylation of histone variant H2A.X (γH2AX) in all cell lines exposed to IR in the presence or absence of ceralasertib treatment (**Fig 4C**). Importantly, combination activity between olaparib and ceralasertib was also observed *in vivo*, with a significant increase in TGI in Δexon11 BRCA1 BTR cell xenografts compared to either monotherapy (**Fig 4D**).

Collectively, these data show increased activity *in vitro* and *in vivo* of the combination of olaparib and ceralasertib in Δexon11 BRCA1 BTR cells, which can be explained by the requirement of ATR activity for RAD51 foci formation in response to DNA damage in this model of BRCA1 Δexon11 hypomorph expression.

## Discussion

Although there are literature examples linking the expression of BRCA1 hypomorphs with resistance to chemotherapy and PARPi, both *in vitro* and *in vivo*, our work provides the most detailed analyses of BRCA1 hypomorphic variants to date. Through the generation of an allelic series including clinically relevant point mutations and deletions, we uncover that the hypomorphic nature of the described Δexon11 variants may reside in their inability to recapitulate the full extent of BRCA1 protein-protein interactions, even under overexpression conditions. Importantly, we also describe a clear dose-response behaviour between BRCA1 Δexon11 protein expression levels and response to PARPi, which may help explain the apparent lack of correlation between the ability of BRCA1m PDX models expressing BRCA1 Δexon11 hypomorphs to form BRCA1 and RAD51 foci, and their response to PARPi (13,28). In addition, our analyses point towards the actual length of the deletion inside the exon 11 of *BRCA1*, rather than any specific regions encoded within, as the main determinant of the hypomorphic nature of these variants. This could explain why, despite the *BRCA1* exon 11 region being the most flexible at accommodating large deletions as secondary reversion mutations potentially driving resistance to chemotherapy and PARPi in patients, deletions resulting in loss of more than 300 amino acids inside the exon 11 sequence are extremely rare events (29).

Our analyses also demonstrate that mutations or deletions blocking the ability of BRCA1 to interact with its partners through the RING, coiled coil or BRCT domains without overtly impacting protein stability all have a drastic effect on BRCA1 function at preventing heightened sensitivity to carboplatin and olaparib. In that regard, only overexpression of the Δexon11 version of BRCA1 generated resistance to olaparib with an IC50 value (approximately 1 μM) above the minimal free concentration of olaparib (approximately 300 nM) in plasma of patients on the established monotherapy dose of 300 mg twice daily (30). Expression of RING and BRCT mutant versions of BRCA1 has been linked to chemotherapy and PARPi resistance in genetically engineered mouse models of BRCA1 deficiency (8) and in PDX models (10,15). Our data are in accordance with previous reports showing that overexpression of RING or BRCT mutant BRCA1 proteins caused modest resistance to chemotherapy and PARPi both *in vitro* and *in vivo* (9,31). Given the clear hypomorphic character of these mutants, we hypothesize that additional mutations and/or DNA repair pathway adaptations are required in tumours for these RING and BRCT hypomorphs to drive clinically meaningful therapy resistance (10,31), making this an area where further research is required.

Our data also highlight that the hypomorphic nature of BRCA1 Δexon11 can be explained, at least in part, by its inability, even at high overexpression levels, to fully complement the deficiency in RAD51 foci formation caused by BRCA1 loss. This can be linked to our observation of an impaired capacity of BRCA1 Δexon11 to interact with PALB2, an interaction that is essential to bring the PALB2-BRCA2-RAD51 complex to sites of DNA damage (5). Importantly, we uncover a specific reliance of our MDAMB436-BTR BRCA1 Δexon11 system on the activity of the DNA damage checkpoint kinase ATR to effectively recruit RAD51 to DNA damage foci. This vulnerability could potentially explain the increased combination activity of ceralasertib plus olaparib in BRCA1 Δexon11 cells *in vitro*, which translated in significant improved efficacy for the combination over either monotherapy *in vivo*, as has been observed in PDX models expressing the BRCA1 Δexon11 hypomorph (32). As the combination of ceralasertib and olaparib is being tested in the clinic (33), our work provides a supportive mechanistic platform to help explain the benefit of this combination in patients with tumours harbouring mutations in the exon 11 of *BRCA1*.

## Materials and methods

### Cell lines and chemicals

HEK-293T (CRL-3216) and MDAMB436 (HTB-130) cells were sourced from ATCC and grown in DMEM (Gibco) or RPMI 1640 (Sigma) respectively, both supplemented with 10% foetal bovine serum (FBS) (Sigma) and 1X GlutaMAX (Gibco). Cells were maintained at 37°C in 5% CO_2._ Olaparib, carboplatin, ceralasertib and adavosertib were made by AstraZeneca and were dosed in DMSO (DMSO normalised across conditions). Tetracycline and doxycycline were sourced from Thermo and Sigma, respectively.

### Constructs

Plasmids containing the human *BRCA1* (NM_007294) and *RAD51B* (NM_133509) open reading frames (ORFs) were obtained from Genscript (Ohu18572D) and OriGene (RC206457L3) respectively. BRCA1 deletion and mutant constructs were generated using the Q5 Site-directed mutagenesis kit (New England Biolabs) and custom primers, sequences confirmed by Sanger sequencing.

### Transient transfection

HEK-293T cells were transfected using a 3.5:1 ratio of FuGENE HD (Promega) to DNA, following manufacturer’s instructions. Cells were lysed for protein after 48/72 hours and samples were processed for western blotting or co-immunoprecipitation.

### Lentivirus generation

HEK-293T cells were transfected with gene of interest lentiviral expression plasmid, psPAX2 (lentiviral packaging plasmid) and mMD2.G VSV-G (lentiviral envelope plasmid) using Lipofectamine LTX (Thermo). Media was changed after 6 hours. 72 hours post-transfection, lentivirus-containing media was harvested, syringe-filtered (0.45 μM) and stored at −80°C.

For *BRCA1*, prior to lentivirus generation, ORFs were cloned into a tetracycline-regulated lentiviral expression vector from the ViraPower™ HiPerform™ T-REx™ Gateway™ Vector Kit (Thermo). This kit also contained a control lentiviral vector for overexpression (LacZ plasmid) and a lentiviral vector for constitutive expression of the tetracycline-repressor protein.

### Cell line generation

MDAMB436 overexpression cell lines were generated by lentiviral transduction. Lentivirus was added to parental cells along with 8 μg/ml polybrene (Millipore) and media was replaced after 24 hours. After a further 24/48 hours, antibiotic was added to select for successfully transduced cells. 1 μg/ml puromycin (Gibco) was used to select for cells infected with the RAD51B lentivirus; 500 μg/ml G418 (Thermo) was used for tetracycline-repressor protein and 12 μg/ml blasticidin (Thermo) was used for BRCA1. Cells were then expanded in the presence of antibiotic and used for the experiments described here. For the MDAMB426-B-TR + BRCA1 cells, tetracycline/doxycycline was added to cells to induce BRCA1 gene expression (1 μg/ml unless otherwise stated).

### (Co-)immunoprecipitation

Cells were lysed for protein in immunoprecipitation lysis buffer (300 mM NaCl, 1 mM EDTA, 20 mM Tris-HCL, 0.5% IGEPAL, 10% glycerol) supplemented with protease inhibitor (Roche). Protein lysates of equal concentration were incubated with Anti-FLAG (Sigma F2426) or Protein A control (Sigma P6486) affinity gel at 4°C for at least 4 hours (up to overnight). Flow through lysates were collected, after which beads were washed with lysis buffer (5 minutes at 4°C, 3 times) and bound proteins were eluted in NuPAGE ™ LDS sample buffer (Invitrogen NP0007) supplemented with reducing agent (Invitrogen NP0009) by heating to 95°C for 5 minutes. Samples were then processed for western blotting.

### Western blotting

Protein lysates were prepared in Laemmli buffer, RIPA buffer (Thermo) (supplemented with phosphatase inhibitor, protease inhibitor, 1% Triton X-100 and 50 U/ml benzonase) or were taken for co-immunoprecipitation experiments. Lysates were normalised to equal protein concentration and mixed with NuPAGE™ LDS sample buffer (Invitrogen NP0007) supplemented with reducing agent (Invitrogen NP0009), before heating to 95°C for 10 minutes. Sodium dodecyl sulfate–polyacrylamide gel electrophoresis was performed by loading samples into NuPAGE™ 4-12% Bis-Tris protein gels and using NuPAGE™ MOPS SDS running buffer (Invitrogen), running at 140V until sample buffer reached the bottom of the gel. Wet transfer onto nitrocellulose (Invitrogen) was performed overnight at 30V, 4°C, followed by blocking with non-fat milk (5% Marvel in TBS, 0.05% Tween). Membranes were incubated in primary antibodies diluted in blocking buffer at 4°C overnight, followed by washing in TBS, 0.05% Tween. Next, membranes were incubated in horseradish peroxidase (HRP)-conjugated secondary antibodies diluted in blocking buffer at room temperature for 1 hour and washed again. Bands were visualised using ECL reagent (Thermo) and imaged using the G:BOX system (Syngene) or X-ray film (Amersham Hyperfilm, GE Healthcare). Band intensity was quantified using ImageJ.

### Colony forming assay

Cells were dosed with olaparib using the HP D300 digital dispenser (Tecan) and cultured for 14 days, after which they were fixed and stained with Blue-G-250 Brilliant Blue (Sigma B8522-1EA) reconstituted in 25% methanol, 5% acetic acid. Once dry, plates were imaged using the GelCount system (Oxford Optronix) and colony density was measured using ImageJ. Curves were plotted using GraphPad Prism and non-linear regression was used to calculate IC50s. Statistical analysis was performed using a One-Way ANOVA with Holm-Sidak post hoc testing.

**Table 1.** Antibodies used for western blotting.

| Target | Host Species | Catalogue Number | Provider | Working Dilution (WB) |
| --- | --- | --- | --- | --- |
| Anti-Mouse IgG HRP | Goat | 7076 | Cell Signaling Tech | 1:2000 – 1:3000 |
| Anti-Rabbit IgG HRP | Goat | 7074 | Cell Signaling Tech | 1:2000 – 1:3000 |
| BARD1 | Rabbit | ab226854 | Abcam | 1:1000 |
| BRCA1 | Mouse | OP92 | Merck | 1:500 – 1:1000 |
| BRCA1 | Rabbit | 07-434 | Merck | 1:500 – 1:1000 |
| BRIP1 | Rabbit | 4578 | Cell Signaling Tech | 1:1000 |
| CTIP (RBBP8) | Rabbit | 9201 | Cell Signaling Tech | 1:1000 |
| GAPDH | Rabbit | 2118 | Cell Signaling Tech | 1:1000 |
| PALB2 | Rabbit | A301-246A | Bethyl Laboratories | 1:1000 |
| RAD51B | Rabbit | GTX40242 | Genetex | 1:1000 |
| Tet Repressor | Mouse | TET01 | MoBiTec | 1:1000 |
| Vinculin | Mouse | V9131 | Sigma-Aldrich | 1:2000 – 1:5000 |

### Survival assays

Cells were dosed with various compounds as single agents using the HP D300 (Tecan) and cultured for 14 days, after which cell viability was assessed using Celltiter-Glo 2 (Promega). Curves were plotted using GraphPad Prism and non-linear regression was used to calculate IC50s. Statistical analysis was performed using a One-Way ANOVA with Holm-Sidak post hoc testing.

For combination assays, cells were dosed using the HP Echo 555 (Labcyte) and cultured for 10 days, after which cell viability was assessed using Celltiter-Glo 2 (Promega). Data were analysed using the Genedata Screener platform, including generating HSA scores for combination activity.

### RT-qPCR

RNA was extracted from *in vitro* cultures using the RNeasy Plus kit (Qiagen) following manufacturer’s instructions. RNA extraction and purification from tumours was performed using the RNeasy 96 QIAcube kit (QIAGEN, #74171). Next, cDNA was synthesised using the SuperScript IV VILO kit (Thermo) following manufacturer’s instructions. RT-qPCR was then performed on the QuantStudio™ 6 Pro (Thermo) using custom primers designed to amplify exogenous FLAG-tagged BRCA1 specifically (or GAPDH as a housekeeping control) and the PowerTrack™ SYBR Green Master Mix (Thermo). Data were analysed using the delta delta Cq method and plotted in GraphPad Prism.

**Table 2.**
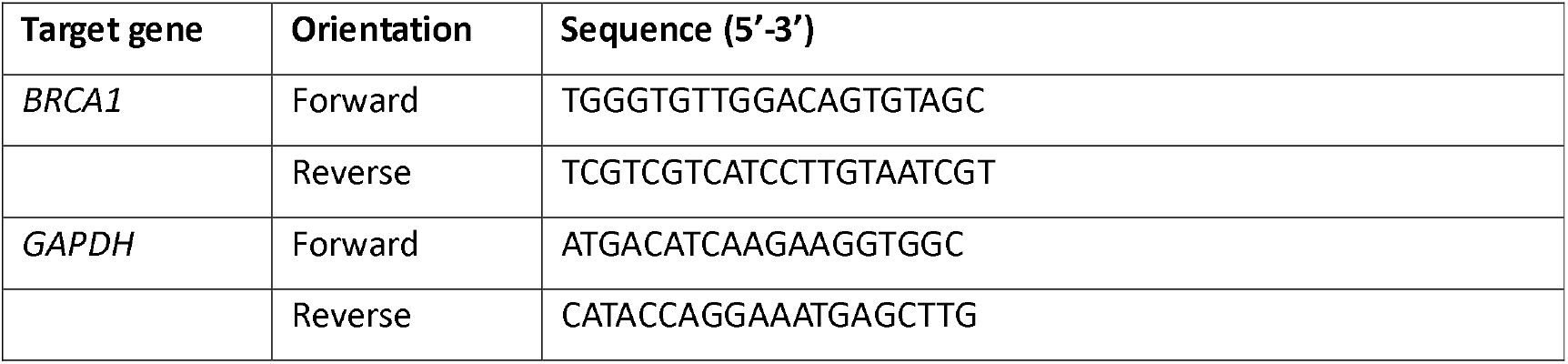
Oligonucleotides used for RT-qPCR.

### Animal studies

All the *in vivo* experimental protocols were monitored and approved by AstraZeneca Animal Welfare and Ethical Review Body, in compliance with guidelines specified by the UK Home Office Animals (Scientific Procedures) Act 1986 and AstraZeneca Global Bioethics policy or Institutional Animal Care and Use Committee (IACUC) guidelines. The Experimental work is outlined in project license PP3292652, which has gone through the AstraZeneca Ethical Review Process. Mice were maintained in a controlled, specific pathogen-free environment at 20°C to 25°C, 40% to 70% humidity, and a 12-hour light-to-dark cycle. Mice were allowed access to food and water *ad libitum* and euthanized at appropriate humane endpoint.

To induce BRCA1 gene expression, doxycycline + sucrose (stock 6.7mg/ml + 135mg/ml) was dissolved in drinking water to reach a final concentration of 0.5mg/ml + 16mg/ml. Water pouch replacement was performed regularly every 4 days, to prevent degradation and contamination.

MDA-MB-436, MDAMB426-B-TR + FL BRCA1 or MDAMB426-B-TR + BRCA1 Δexon11 cells (5×10^6^with 50% Matrigel (Corning)) were implanted subcutaneously to SCID (C.B-17/IcrHsd-*Prkdc* ^*scid*^) female mice (Envigo, UK). Animal body weight and tumour condition were monitored throughout the study. Tumour length and width were measured by calliper, and tumour volume (TV) was calculated using the formula volume = (length × width^2^)^*^π/6. Mice were randomized into treatment groups when mean TV reached approximately 0.2 cm^3^.

Tumour growth inhibition (TGI) from the start of treatment was assessed by comparison of the mean change in TV of the control and treated groups and represented as per cent TGI (when TV ≥ starting TV) or tumour regression (TR, when TV < starting TV). Statistical significance was evaluated using a one-tailed t test.

Olaparib was formulated in 10% DMSO, 30% kleptose and dosed continuously; ceralasertib was formulated in 10% DMSO, 40% propylene glycol and dosed with schedule 3 times/weekly, BID. Mice received treatment for a maximum of 42 days following randomization.

### Immunofluorescence

Cells were seeded in 96-well plates using media containing 1 μg/mL doxycycline to induce BRCA1 expression. The following day, cells were treated with 0.3 μM ceralasertib for 24 hours. Subsequently, cells were exposed to 5 Gy of ionizing radiation (IR) using a high-voltage X-ray generator (Faxitron X-Ray Corporation). Four hours post-IR, cells were fixed with 4% paraformaldehyde for 15 minutes at room temperature (RT), followed by permeabilisation with 1× PBS containing 0.1% Triton X-100 for 10 minutes at RT. Blocking was performed using 0.5% BSA and 0.2% gelatin from cold-water fish skin (Sigma) in 1× PBS for 1 hour at RT.

Cells were then incubated overnight at 4°C with primary antibodies, followed by incubation at RT with Alexa Fluor-conjugated secondary antibodies and DAPI (Sigma, 1 μg/mL) for 1.5 hours.

Image acquisition was carried out using an Olympus ScanR microscope with a 40× objective to analyse DNA damage-induced foci. Cell cycle distribution was determined based on DAPI staining intensity, and foci were quantified using the “spots detector module” in cells at the S–G2 phases.

**Table 3.** Antibodies used for immunofluorescence experiments.

| Target | Host Species | Catalogue Number | Provider | Working Dilution (IF) |
| --- | --- | --- | --- | --- |
| Alexa Fluor 488 | Goat (anti-mouse) | A-11029 | Thermo Fisher Scientific | 1:2000 |
| Alexa Fluor 594 | Goat (anti-rabbit) | A-11037 | Thermo Fisher Scientific | 1:2000 |
| γH2AX | Mouse | 05-636 | Millipore (Sigma/Merck) | 1:5000 |
| RAD51 | Rabbit | 70-001 | Bioacademia | 1:7000 |

## Acknowledgements

The authors would like to thank Dr Ross Hill and Dr Domenic Pilger for critical reading of the manuscript.

## Supplementary figure legends

**Supplementary Figure S1.**
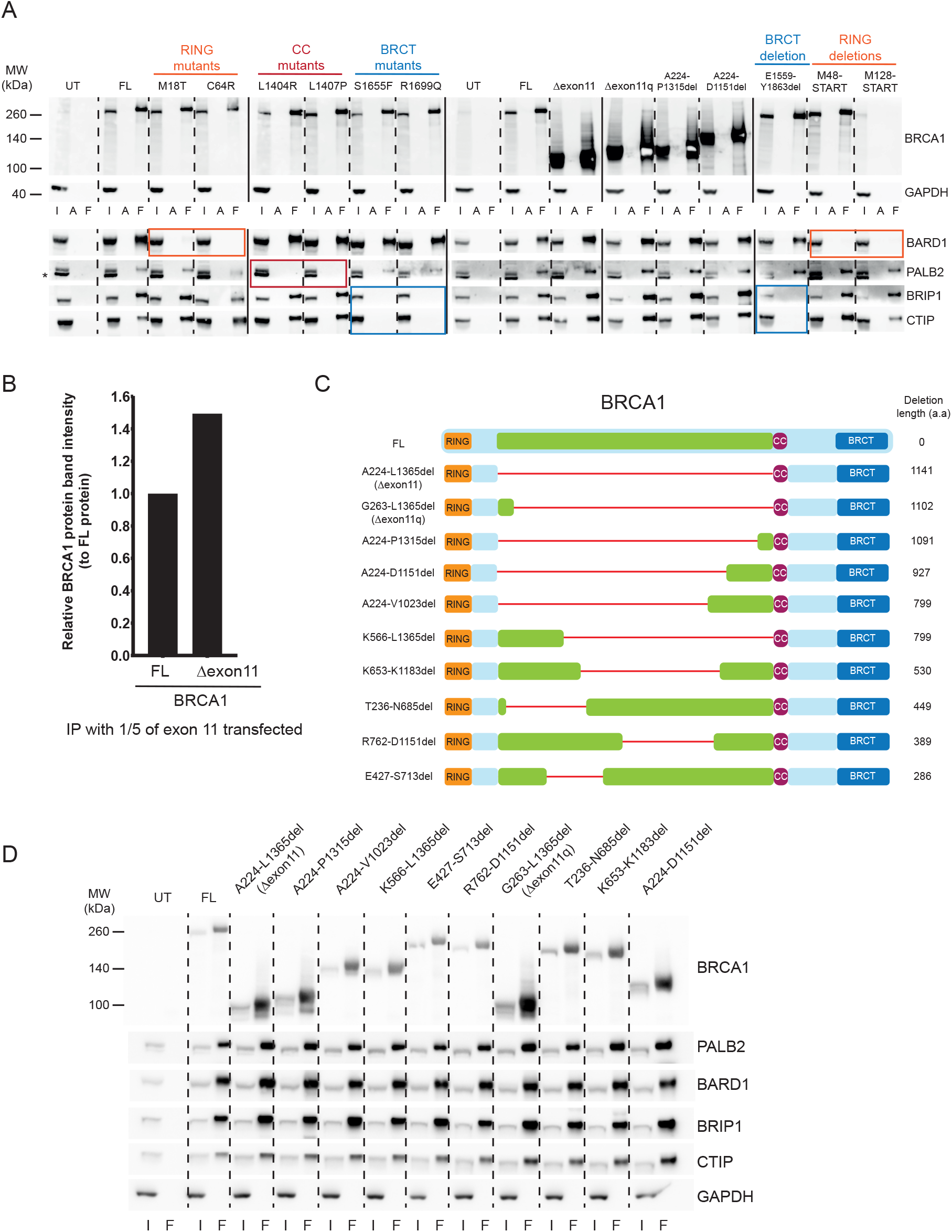
**A**) Co-immunoprecipitation experiments in HEK293T cells with the different forms of BRCA1 used in this study. UT: unstransfected; FL: full length; I: input; A: pull down with protein A beads; F: pull down with FLAG beads. **B**) Quantification of the relative BRCA1 protein pull down normalized to the BRCA1 full length (FL) protein, related to Figure 1D. **C**) Schematic of the BRCA1 exon 11 protein deletions analysed in this study. **D**) Co-immunoprecipitation experiments in HEK293T cells with the different exon 11 deletions of BRCA1 used in this study. UT: unstransfected; FL: full length; I: input; F: pull down with FLAG beads.

**Supplementary Figure S2.**
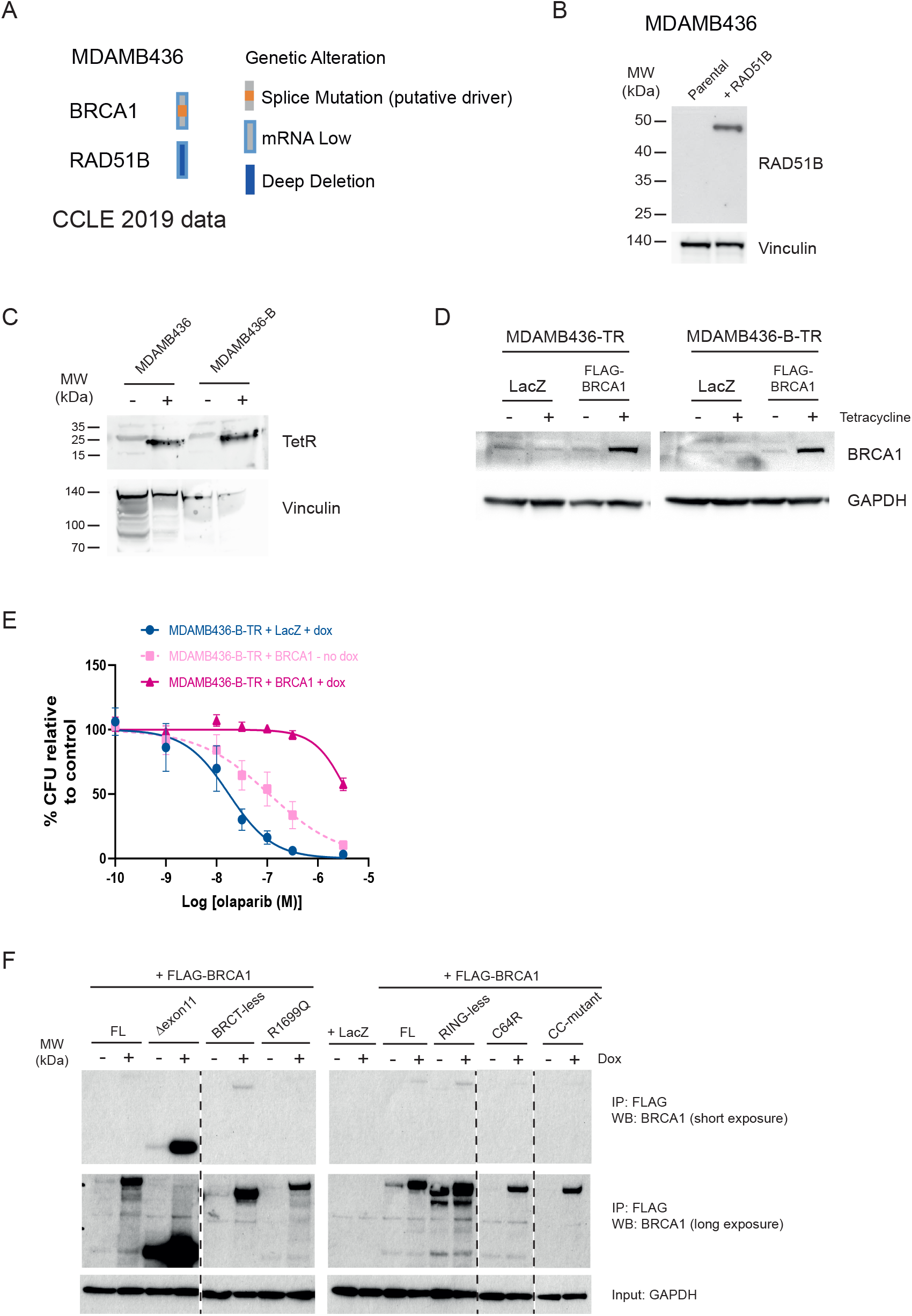
**A**) Genetic alterations in the MDAMB436 cell line, as described in the Cancer Cell Line Encyclopaedia data update of 2019. **B**) Western blot showing expression of RAD51B only in MDAMB436 cells stably transfected with a RAD51B expression construct. Vinculin was used as loading control. **C**) Western blot showing expression of the tetracycline repressor (TetR) only in MDAMB436 or MDAMB436-B (+ RAD51B) cells transfected with the TetR construct (+). Vinculin was used as loading control. **D**) Western blot showing expression of FLAG-tagged BRCA1 in the presence of tetracycline induction in MDAMB436-TR (+ TetR) or MDAMB436-B-TR (+ RAD51B + TetR) cells. GAPDH was used as loading control. **E**) Dose-response curves of olaparib in colony formation assays in tetracycline-repressor (TR) expressing MDAMB436-B (+ RAD51B) cells with (+ BRCA1) or without (+ LacZ) BRCA1 complementation, cultured in the presence or absence of doxycycline (+/-dox). **F**) Western blot showing immunoprecipitation of FLAG-tagged BRCA1 constructs (or LacZ control) stably transfected in MDAMB436-B-TR cells, in the presence or absence of doxycycline induction. FL: full length. GAPDH was used as input control.

## References

1. Prados-Carvajal R, Irving E, Lukashchuk N, Forment JV.x Preventing and Overcoming Resistance to PARP Inhibitors: A Focus on the Clinical Landscape. Cancers (Basel) 2022;14:44

2. Krais JJ, Johnson N. BRCA1 Mutations in Cancer: Coordinating Deficiencies in Homologous Recombination with Tumorigenesis. Cancer Research 2020;80:4601–9

3. Densham RM, Garvin AJ, Stone HR, Strachan J, Baldock RA, Daza-Martin M, et al. Human BRCA1-BARD1 ubiquitin ligase activity counteracts chromatin barriers to DNA resection. Nat Struct Mol Biol 2016;23:647–55

4. Scully R, Chen J, Plug A, Xiao Y, Weaver D, Feunteun J, et al. Association of BRCA1 with Rad51 in Mitotic and Meiotic Cells. Cell 1997;88:265–75

5. Anantha RW, Simhadri S, Foo TK, Miao S, Liu J, Shen Z, et al. Functional and mutational landscapes of BRCA1 for homology-directed repair and therapy resistance. Elife 2017;6

6. Rebbeck TR, Mitra N, Wan F, Sinilnikova OM, Healey S, McGuffog L, et al. Association of type and location of BRCA1 and BRCA2 mutations with risk of breast and ovarian cancer. JAMA 2015;313:1347–61

7. Drost R, Bouwman P, Rottenberg S, Boon U, Schut E, Klarenbeek S, et al. BRCA1 RING function is essential for tumor suppression but dispensable for therapy resistance. Cancer Cell 2011;20:797–809

8. Drost R, Dhillon KK, van der Gulden H, van der Heijden I, Brandsma I, Cruz C, et al. BRCA1185delAG tumors may acquire therapy resistance through expression of RING-less BRCA1. J Clin Invest 2016;126:2903–18

9. Wang Y, Krais JJ, Bernhardy AJ, Nicolas E, Cai KQ, Harrell MI, et al. RING domain–deficient BRCA1 promotes PARP inhibitor and platinum resistance. The Journal of Clinical Investigation 2016;126:3145–57

10. Pellegrino B, Herencia-Ropero A, Llop-Guevara A, Pedretti F, Moles-Fernández A, Viaplana C, et al. Preclinical In Vivo Validation of the RAD51 Test for Identification of Homologous Recombination-Deficient Tumors and Patient Stratification. Cancer Research 2022;82:1646–57

11. Tammaro C, Raponi M, Wilson David I, Baralle D. BRCA1 exon 11 alternative splicing, multiple functions and the association with cancer. Biochemical Society Transactions 2012;40:768–72

12. Wang Y, Bernhardy AJ, Cruz C, Krais JJ, Nacson J, Nicolas E, et al. The BRCA1-Delta11q Alternative Splice Isoform Bypasses Germline Mutations and Promotes Therapeutic Resistance to PARP Inhibition and Cisplatin. Cancer Res 2016;76:2778–90

13. Cruz C, Castroviejo-Bermejo M, Gutiérrez-Enríquez S, Llop-Guevara A, Ibrahim YH, Gris-Oliver A, et al. RAD51 foci as a functional biomarker of homologous recombination repair and PARP inhibitor resistance in germline BRCA-mutated breast cancer. Annals of Oncology 2018:mdy099-mdy

14. Johnson N, Johnson SF, Yao W, Li Y-C, Choi Y-E, Bernhardy AJ, et al. Stabilization of mutant BRCA1 protein confers PARP inhibitor and platinum resistance. Proceedings of the National Academy of Sciences 2013;110:17041–6

15. Wang Y, Bernhardy AJ, Nacson J, Krais JJ, Tan Y-F, Nicolas E, et al. BRCA1 intronic Alu elements drive gene rearrangements and PARP inhibitor resistance. Nature Communications 2019;10:5661

16. Bouwman P, van der Gulden H, van der Heijden I, Drost R, Klijn CN, Prasetyanti P, et al. A high-throughput functional complementation assay for classification of BRCA1 missense variants. Cancer Discov 2013;3:1142–55

17. Nacson J, Di Marcantonio D, Wang Y, Bernhardy AJ, Clausen E, Hua X, et al. BRCA1 Mutational Complementation Induces Synthetic Viability. Molecular Cell 2020

18. Lin KK, Harrell MI, Oza AM, Oaknin A, Ray-Coquard I, Tinker AV, et al. BRCA Reversion Mutations in Circulating Tumor DNA Predict Primary and Acquired Resistance to the PARP Inhibitor Rucaparib in High-Grade Ovarian Carcinoma. Cancer Discov 2019;9:210–9

19. Kondrashova O, Nguyen M, Shield-Artin K, Tinker AV, Teng NNH, Harrell MI, et al. Secondary Somatic Mutations Restoring RAD51C and RAD51D Associated with Acquired Resistance to the PARP Inhibitor Rucaparib in High-Grade Ovarian Carcinoma. Cancer Discovery 2017;7:984–98

20. Elstrodt F, Hollestelle A, Nagel JHA, Gorin M, Wasielewski M, van den Ouweland A, et al. BRCA1 Mutation Analysis of 41 Human Breast Cancer Cell Lines Reveals Three New Deleterious Mutants. Cancer Res 2006;66:41–5

21. Illuzzi G, Staniszewska AD, Gill SJ, Pike A, McWilliams L, Critchlow SE, et al. Preclinical Characterization of AZD5305, A Next-Generation, Highly Selective PARP1 Inhibitor and Trapper. Clinical Cancer Research 2022;28:4724–36

22. Ghandi M, Huang FW, Jané-Valbuena J, Kryukov GV, Lo CC, McDonald ER, et al. Next-generation characterization of the Cancer Cell Line Encyclopedia. Nature 2019;569:503–8

23. Jamal K, Galbiati A, Armenia J, Illuzzi G, Hall J, Bentouati S, et al. Drug–gene Interaction Screens Coupled to Tumor Data Analyses Identify the Most Clinically Relevant Cancer Vulnerabilities Driving Sensitivity to PARP Inhibition. Cancer Research Communications 2022;2:1244–54

24. Veneris JT, Matulonis UA, Liu JF, Konstantinopoulos PA. Choosing wisely: Selecting PARP inhibitor combinations to promote anti-tumor immune responses beyond BRCA mutations. Gynecologic Oncology 2020;156:488–97

25. Wilson Z, Odedra R, Wallez Y, Wijnhoven PWG, Hughes AM, Gerrard J, et al. ATR Inhibitor AZD6738 (Ceralasertib) Exerts Antitumor Activity as a Monotherapy and in Combination with Chemotherapy and the PARP Inhibitor Olaparib. Cancer Research 2022;82:1140–52

26. Hirai H, Iwasawa Y, Okada M, Arai T, Nishibata T, Kobayashi M, et al. Small-molecule inhibition of Wee1 kinase by MK-1775 selectively sensitizes p53-deficient tumor cells to DNA-damaging agents. Mol Cancer Ther 2009;8:2992–3000

27. Berenbaum MC. What is synergy? Pharmacological Reviews 1989;41:93–141

28. Castroviejo-Bermejo M, Cruz C, Llop-Guevara A, Gutiérrez-Enríquez S, Ducy M, Ibrahim YH, et al. A RAD51 assay feasible in routine tumor samples calls PARP inhibitor response beyond BRCA mutation. EMBO Molecular Medicine 2018

29. Tobalina L, Armenia J, Irving E, O’Connor MJ, Forment JV. A meta-analysis of reversion mutations in BRCA genes identifies signatures of DNA end-joining repair mechanisms driving therapy resistance. Annals of Oncology 2021;32:103–12

30. Mateo J, Moreno V, Gupta A, Kaye SB, Dean E, Middleton MR, et al. An Adaptive Study to Determine the Optimal Dose of the Tablet Formulation of the PARP Inhibitor Olaparib. Targeted Oncology 2016;11:401–15

31. Nacson J, Krais JJ, Bernhardy AJ, Clausen E, Feng W, Wang Y, et al. BRCA1 Mutation-Specific Responses to 53BP1 Loss-Induced Homologous Recombination and PARP Inhibitor Resistance. Cell Rep 2018;24:3513–27 e7

32. Serra V, Wang AT, Castroviejo-Bermejo M, Polanska UM, Palafox M, Herencia-Ropero A, et al. Identification of a Molecularly-Defined Subset of Breast and Ovarian Cancer Models that Respond to WEE1 or ATR Inhibition, Overcoming PARP Inhibitor Resistance. Clinical Cancer Research 2022;28:4536–50

33. Wethington SL, Shah PD, Martin L, Tanyi JL, Latif N, Morgan M, et al. Combination ATR (ceralasertib) and PARP (olaparib) Inhibitor (CAPRI) Trial in Acquired PARP Inhibitor-Resistant Homologous Recombination-Deficient Ovarian Cancer. Clin Cancer Res 2023;29:2800–7

